# Did we get sensorimotor adaptation wrong? Implicit adaptation as direct policy updating rather than forward-model-based learning

**DOI:** 10.1101/2020.01.22.914473

**Authors:** Alkis M. Hadjiosif, John W. Krakauer, Adrian M. Haith

## Abstract

The human motor system can rapidly adapt its motor output in response to errors. The prevailing theory of this process posits that the motor system adapts an internal forward model that predicts the consequences of outgoing motor commands, and that this forward model is then used to guide selection of motor output. However, although there is clear evidence for the existence of adaptive forward models to help track the state of the body, there is no real evidence that such models influence the selection of motor output. A possible alternative to the forward-model-based theory of adaptation is that motor output could be directly adjusted by movement errors (“direct policy learning”), in parallel with but independent of any updates to a predictive forward model. Here, we show evidence for this latter theory based on the properties of implicit adaptation under mirror-reversed visual feedback. We show that implicit adaptation still occurs under this extreme perturbation but acts in an inappropriate direction, following a pattern consistent with direct policy learning but not forward-model-based learning. We suggest that the forward-model-based theory of adaptation needs to be re-examined and that direct policy learning is a more plausible mechanism of implicit adaptation.

## Introduction

When we make errors in our movements, the motor system automatically adapts its output in the next movement in order to reduce the error (Krakauer et al., 2019; Shadmehr et al., 2010). This capacity, referred to as *adaptation*, has been demonstrated across a wide variety of behaviors including eye movements (Kojima et al., 2004; Wong & Shelhamer, 2011), reaching movements (Hadjiosif & Smith, 2015; Krakauer et al., 2000, 2005; Shadmehr & Mussa-Ivaldi, 1994; Smith et al., 2006), locomotion (Jayaram et al., 2012; Malone et al., 2011), and speech (Parrell et al., 2017).

The prevailing theory of adaptation is that it is driven by updating of an internal forward model (Figure 1A) – a network within the brain that predicts the consequences of outgoing motor commands (Bastian, 2006; Bhushan & Shadmehr, 1999; Flanagan et al., 2003; A. M. Haith & Krakauer, 2013; Krakauer & Mazzoni, 2011; Krakauer & Shadmehr, 2006; Shadmehr et al., 2010). There is strong behavioral (Bhanpuri et al., 2013; Izawa & Shadmehr, 2011; R. Chris Miall et al., 2007; Synofzik et al., 2008; Wagner & Smith, 2008) and neurophysiological (Ebner & Pasalar, 2008) evidence for the existence of forward models and that these forward models are updated when the consequences of our motor commands are altered by external perturbations (Ebner & Pasalar, 2008; Izawa & Shadmehr, 2011; McNamee & Wolpert, 2019; Synofzik et al., 2008). However, such changes in a forward model do not inherently prescribe how motor output should change in the future; they simply allow the motor system to anticipate the consequences of a given motor command. In order to drive changes in behavior, an adapted forward model must somehow influence the control policy that determines which actions are selected to achieve a particular movement goal. It is unclear exactly how this might occur. Nevertheless, a broad consensus seems to have emerged that forward-model-based learning of this kind is the mechanism by which our movements are adapted when we experience movement errors (Bastian, 2006; Flanagan et al., 2003; A. M. Haith & Krakauer, 2013; Haruno et al., 2001; Jordan & Rumelhart, 1992; R Christopher Miall & Wolpert, 1996; Shadmehr et al., 2010; Wolpert & Kawato, 1998).

**Figure 1.**
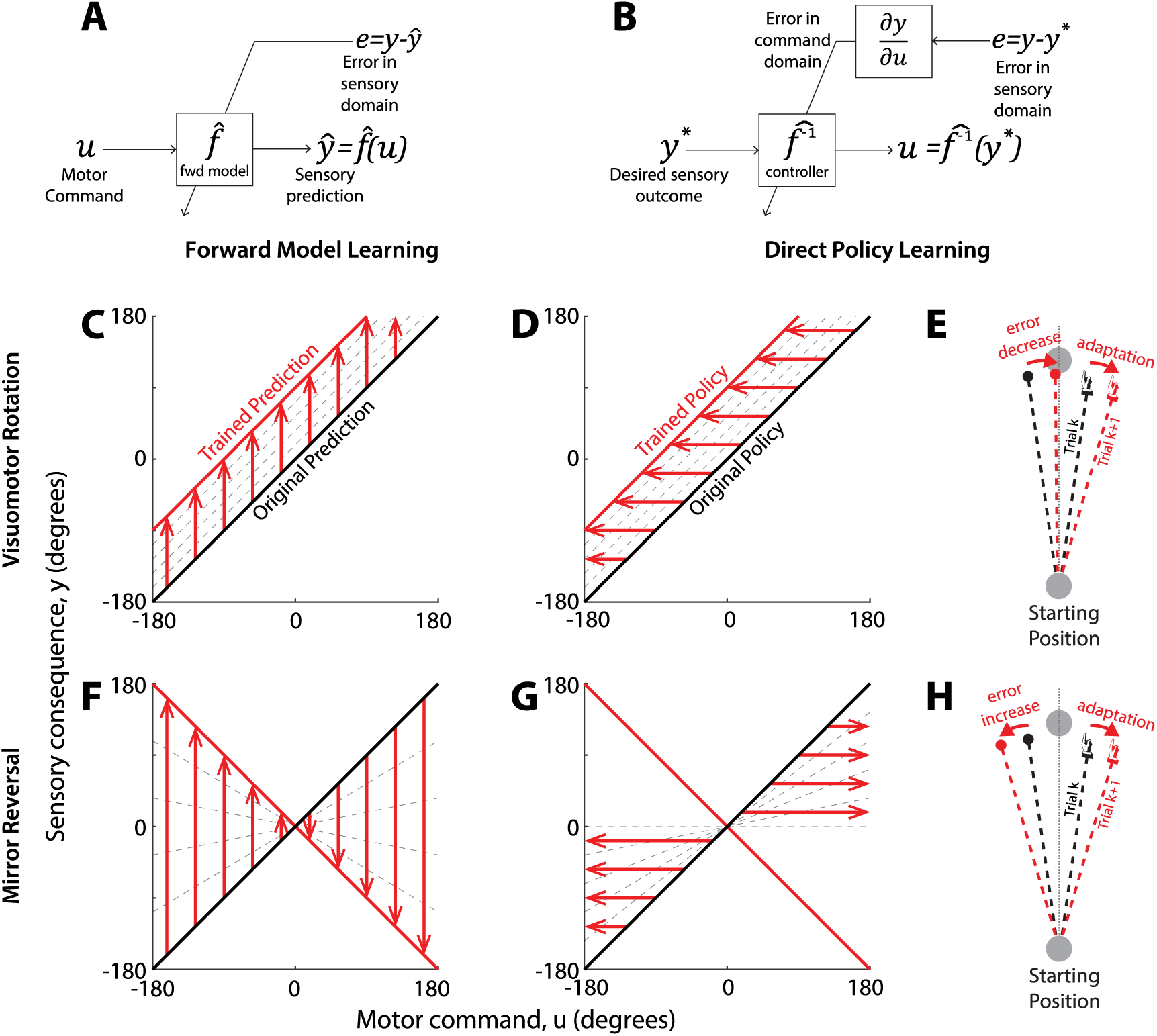
Forward-model-based learning and direct policy learning under visuomotor rotation and mirror reversal. **A:** Forward-model-based learning relies on updating an internal forward model which predicts the sensory consequences (y) of motor commands (u). In the case of simple cursor perturbations, we take u to be the direction the hand moves (motor command) and y to be the direction the cursor will move (sensory outcome). When the predicted sensory outcome (ŷ) differs from the observed one (y), the resulting error can be used to update the forward model without further assumptions. The updated forward model can then be inverted to yield the appropriate motor command. **B: In** direct policy learning, sensory errors are used directly to update the policy (also often called an ‘inverse model’). Here, the policy is a function mapping movement goals (y*), represented in terms of desired sensory outcomes, to appropriate motor commands (u). Sensory errors must be translated to the motor domain, but this mapping depends on knowledge of the plant f (in mathematical terms, knowledge of the sensitivity derivative ∂f/∂u). In practice, direct policy learning must use an assumed mapping (Abdelghani & Tweed, 2010). **C**,**D:** Performance of forward-model-based learning (C) and Direct policy learning (D) under visuomotor rotation. In this example, y = u at baseline (black line), and direct policy updating therefore assumes that e_y_ = e_u_. The red line represents the perturbed mapping (a 90° visuomotor rotation, y = u + 90). Under both models, adaptive changes are appropriate for the shifted visuomotor map: the forward model adjusts to predict the shifted sensory outcomes of efferent motor commands, whereas direct policy updates drive motor output in a direction that reduces error. **E:** Response of direct policy learning to errors resulting from a visuomotor rotation. In response to a leftward error, motor output is shifted rightwards, reducing error in the next trial. **F**,**G:** Forward-model-based learning (F) and Direct policy learning (G) under mirror reversal. Under the mirror-reversal, y=-u, and therefore the relationship between task errors and motor errors is inverted: e_y_=-e_u_. If the sensitivity derivative used for direct policy learning is not updated accordingly, policy updates will occur in the wrong direction, driving motor output further and further away from that necessary to counter the mirror reversal. By contrast, there is no such difficulty associated with learning an updated forward model under the mirror reversal and therefore forward-model-based learning can occur without any issue. **H:** Response of direct policy learning to errors resulting from a mirror reversal. In response to a leftward error, motor output is shifted rightwards, exactly as in (E). Under the mirror reversal, however, this shift leads to an increase in error on the next trial.

An alternative mechanism of adaptation is that errors could simply be used to directly adjust motor output (Figure 1B). For example, if you missed a target to the right, you can simply adjust your motor output to the left next time (Figure 1E), without ever needing to consult a predictive forward model. This much more straightforward mechanism – which we refer to as direct policy learning – has been proposed in the past as a potential mechanism for motor adaptation (Abdelghani et al., 2008; Wolpert et al., 2001), but has become largely overlooked in favor of forward-model-based learning. Both theories, however, generate more or less identical predictions about adaptation in conventional adaptation paradigms; on current evidence it is impossible to distinguish between them.

In order to dissociate between these alternative theories of adaptation, we examine behavior under reversal of visual feedback. Mirror-reversal is a much more drastic perturbation than those that are usually thought to engage implicit adaptation (e.g. visuomotor rotations). Nevertheless, sensory prediction errors still occur under a mirror-reversed visual feedback, particularly for movements close to the mirroring axis, and close examination of behavior has revealed clear signatures of implicit adaptation occurring when people learn to act under a mirror reversal (Lillicrap et al., 2013; Wilterson & Taylor, 2019). In these circumstances, however, implicit adaptation seems to drive motor output in the wrong direction – acting to *increase* rather than decrease errors from one trial to the next. Participants are unable to successfully compensate for the mirror reversal using only implicit adaptation. Instead, they must engage additional, likely explicit, learning mechanisms to achieve successful compensation (Telgen et al., 2014; Wilterson & Taylor, 2019). This fact has led to a supposition that mirror reversal is an unsuitable paradigm for studying implicit adaptation. On the contrary, however, we suggest that understanding the properties of implicit adaptation under mirror reversal – and in particular why it fails to allow compensation – can enable us to dissociate between alternative underlying mechanisms.

In principle, forward-model-based learning ought to have no difficulty compensating for a mirror reversal, even though the perturbation is quite drastic; if the cursor went further to the right than predicted, the forward model should just predict that the cursor will move further to the right in the future. This process will ultimately converge on an accurate forward model regardless of whether it was a rotation or mirror reversal that caused the error. Thus, it is not obvious why adaptation should fail under mirror-reversal if it is based on updating a forward model.

By contrast, the inappropriate adaptation seen under mirror-reversal is expected if adaptation is based on direct policy learning. Direct policy learning requires that an observed error in the *sensory* domain be translated into an error in the *motor* domain, in order to adjust motor output in future movements. This translation necessarily requires an assumption about the relationship between motor output and sensory outcomes (A. Haith & Vijayakumar, 2009; Porrill et al., 2004), i.e. that a leftward visual error should be compensated by reaching further to the right in the next trial (formally, this assumption is termed a *sensitivity derivative* (Abdelghani et al., 2008)). Under a mirror reversal, the contingency between motor output and sensory outcomes becomes flipped and an observed leftward error should now be corrected by reaching further to the left Unless the learning rule is reversed accordingly, error-driven updates will drive behavior in the wrong direction, increasing errors over time rather than reducing them (Figure 1G,H). This pattern of behavior expected under direct policy learning is precisely what is observed under mirror reversal.

The phenomenon of implicit adaptation under mirror-reversed visual feedback thus provides a potential means to dissociate whether adaptation occurs through forward-model-based learning or through direct policy learning. Although existing accounts of implicit learning under mirror-reversal seem consistent with direct policy adaptation and not forward-model-based adaptation, this phenomenon has not been sufficiently well characterized to conclusively support this logic. One important limitation of existing work is that imposing a mirror reversal is likely to engage additional learning processes besides implicit adaptation (Gutierrez-Garralda et al., 2013; Telgen et al., 2014), such as strategic re-aiming (McDougle et al., 2016; Morehead et al., 2015; Wilterson & Taylor, 2019). It is possible that these other processes are responsible for the observed mal-adaptation, rather than implicit recalibration. Furthermore, although a perfectly adapted forward model should in theory enable perfect compensation, an incompletely adapted forward model could also lead to catastrophic failure of forward-model-based learning, as we examine in more detail below. We therefore conducted a series of experiments designed to address these limitations and establish key properties of implicit adaptation under mirror reversal and dissociate whether forward-model-based learning or direct policy learning can account for implicit adaptation.

## Results

### Implicit adaptation amplifies errors at targets along the mirroring axis

In Experiment 1, 12 participants made planar 12cm reaching movements to “shoot” through one of four different targets arranged on the cardinal directions (Figure 2A,B): two targets positioned on the mirroring axis (on-axis targets) and two perpendicular to it (off-axis targets). We began with three blocks of unperturbed movements to familiarize participants with the task and to assess baseline behavior. This was followed by the training phase, whereby we introduced a mirror reversal about the y-axis. To isolate implicit adaptation to this mirror reversal, we instructed participants to aim their *hand* directly through the target, rather than try to guide the mirrored *cursor* through the target, even though the cursor might miss it. This approach has been demonstrated to successfully isolate implicit components of adaptation (Kim et al., 2018; Morehead et al., 2015). All participants but one, who was excluded from further analysis, were successful in following this instruction, as evidenced by their performance at off-axis targets, where they overwhelmingly (98.1±0.5% of off-axis trials, mean±SEM) aimed their hand towards the target, though this caused the cursor to move in the opposite direction.

**Figure 2.**
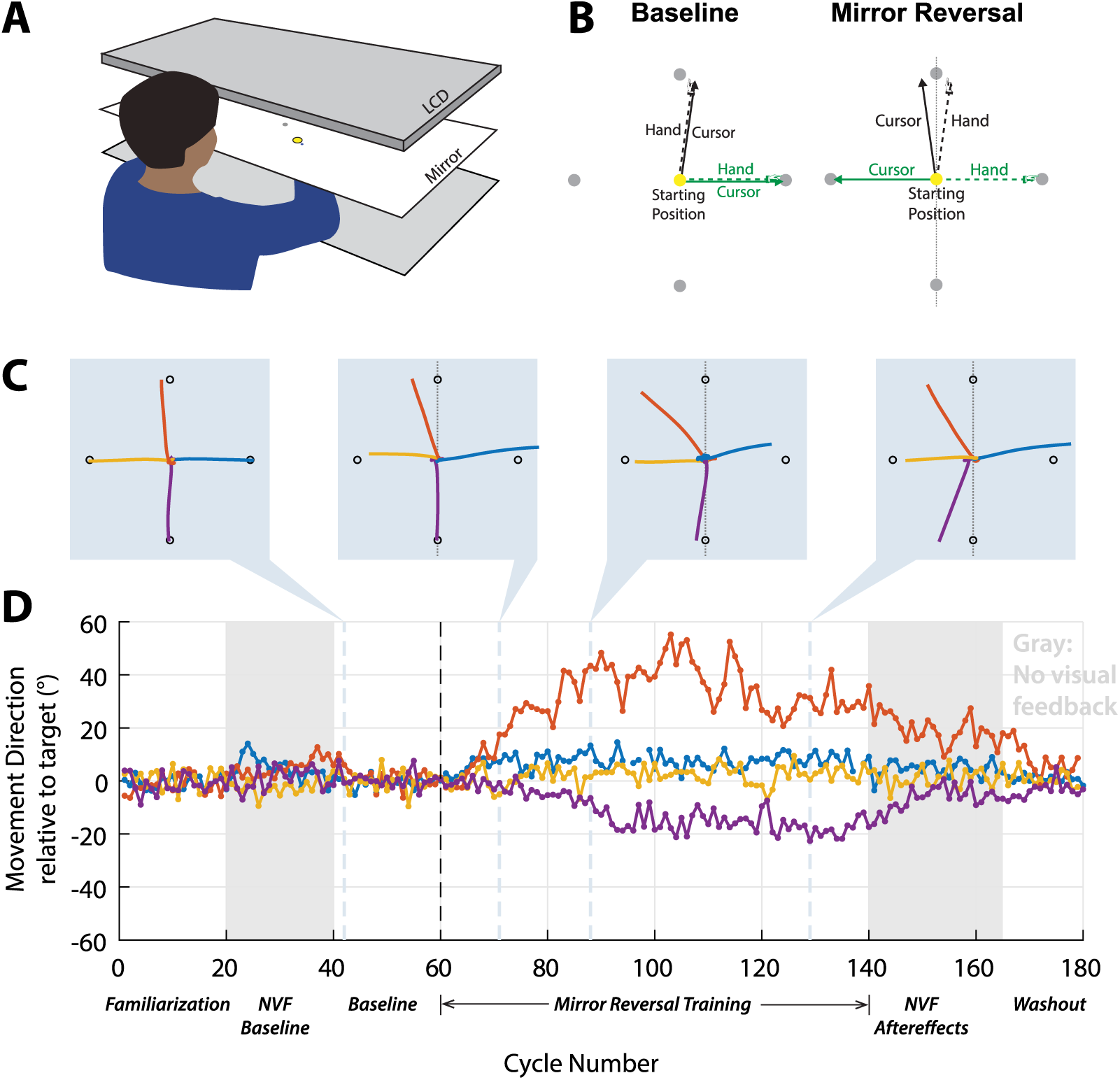
Experiment 1 design and example participant. **A:** Participants made reaching movements on a tabletop. Vision of the hand was occluded and an LCD screen projected a blue cursor which appeared at the level of the table through a mirror. **B:** Participants made center-out shooting movements from a central starting position (yellow circle) through different targets (gray circles). During baseline (left), the cursor followed hand position; during mirror reversal training (right), cursor feedback was reflected about the y-axis (vertical dashed line). **C:** Example hand trajectories from (left to right) baseline, early, mid, and late learning, for one participant. **D:** Movement direction relative to the target for all target directions. There is a clear drift away from the two targets along the mirroring axis (orange and purple), which persists after visual feedback is removed after trial 140 (gray). For clarity, trials to horizontal targets in which the participant moved towards the opposing target are omitted from this plot (3 trials in blue curve).

Figure 2C,D shows the behavior of an example participant at different points during the experiment. After the mirror-reversal was introduced (cycles 61-140) there was little, if any, change in the direction of reaching movements to off-axis targets (yellow and blue). For reaching movements to on-axis targets (orange and purple), however, small initial deviations tended to be amplified from trial to trial, rather than corrected, as illustrated in 2D. These deviations tended to saturate at around 20-30° away from the axis, which is consistent with the limited capacity of the implicit adaptation system identified in visuomotor rotation studies (Bond & Taylor, 2015; Kim et al., 2018; Morehead et al., 2017).

Behavior of this example participant was similar to that observed across the population as a whole (Figure 3A), with small deviations becoming amplified, resulting in a drift away from the mirroring axis (0° in Figure 3A). The direction of this drift varied across participants, directed either clockwise or counterclockwise (negative or positive values, respectively, on the same panel). However, the absolute directional error for reaches to the two on-axis targets increased significantly from baseline to late adaptation (2.5±0.2° in baseline vs. 17.8±1.3° in the last two mirror training blocks (6^th^ and 7^th^), p <10^−6^, shown in Figure 3D). A post-hoc analysis showed that the direction of the initial deviations and subsequent drift generally followed directional biases of baseline hand movements in almost every case (for 10/11 participants for the “up”, 90° target, and 8/11 participants for the “down”, 270° target as shown in Figure 3C). Although participants generally showed a monotonic drift that was either consistently clockwise or counterclockwise relative to the target, a small number of participants (2/11 at the “up” target and 1/11 at the “down” target) reversed the direction of their drift between the first and fourth training block, as illustrated in Figure3A.

**Figure 3.**
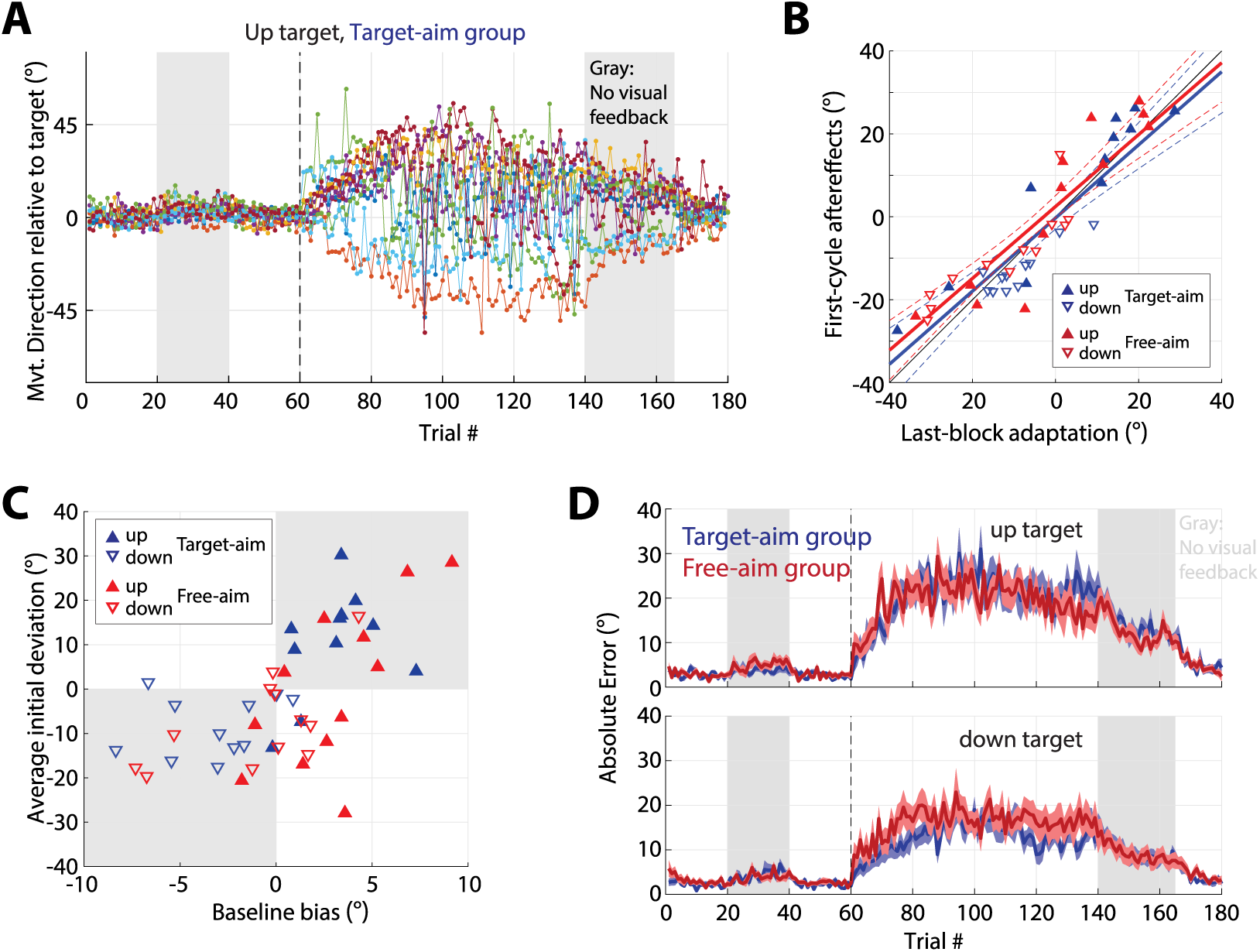
Instability of implicit adaptation under a mirror reversal. **A:** Errors in hand direction for the 90° (“up”) target in Experiment 1. Each curve shows data from one participant. The mirror reversal was imposed during trials 61-140. Gray regions indicate trials during which visual feedback was withheld. The data show participants’ reach directions immediately beginning to drift away from the ideal direction. This instability is consistent with implicit adaptation occurring through direct policy learning but not forward-model-based learning. Removal of visual feedback (grey region) led to clear and persistent aftereffects. **B:** The magnitude of aftereffects in Experiments 1 and 1a matched the level of drift during late exposure to the mirror reversal. Plotted is the amount of aftereffects (first trial after visual feedback removal, trial 141) against the corresponding level of late drift (last 20 trials of preceding block). Upwards-pointing triangles: “up” target; Downwards-pointing triangles: “down” target; Blue: Target-aim group (Experiment 1); Red: Free-aim group (Experiment 1a). Solid lines show linear fits, colored dashed lines indicate the corresponding 95% confidence intervals; black thin solid line indicates the unity line. **C:** The direction of (mal-)adaptive changes during mirror reversal training is predicted by directional biases measured during baseline. Biases measured during the second experiment block (trials 21-40) without visual feedback, show a strong correspondence with the initial mal-adaptive changes measured during the first training block (trials 61-80). The shaded area indicates points where the direction of baseline biases and initial mal-adaptive changes are congruent. **D:** Absolute errors for the “up” (top) and “down” (bottom) targets in for the Target-aim group (Experiment 1, blue) and Free-aim group (Experiment 1a, red). Shading indicates SEM.

Before the 8^th^ (penultimate) block, visual feedback was removed and participants were reminded to keep aiming their hand through the target. The example participant in Figure 2 (gray, trials 141-165) exhibited strong, slowly-decaying aftereffects during this block. This pattern was consistent across the whole group of participants, which showed strong aftereffects during the no-visual-feedback block (absolute reaching angle on the first post-learning cycle: 16.2±0.9° vs. 4.1±0.4° for the no-visual-feedback baseline (block 2), p <10^−7^). Furthermore, these aftereffects closely matched the amount of drift during the last exposure block (last 20 adaptation trials) as shown in blue in Figure 3B (linear fit between subject-averaged reaching angles during the last exposure block and the first aftereffect cycle: slope = 0.88 (95% c.i.: [0.66, 1.10], R^2^ = 0.90, p<10^−5^). These aftereffects persisted throughout the no-visual-feedback block, decaying slowly, and were only completely extinguished once veridical feedback was restored, as shown in Figure 3A. This gradual decay of aftereffects is a key signature of implicit adaptation (Galea et al., 2010; Kitago et al., 2013) and thus confirms that the change in behavior during the exposure blocks was attributable to implicit adaptation.

In summary, this experiment showed that implicit adaptation occurred under mirror reversal, but did not act to reduce performance errors; instead, it amplified initially small errors, driving reaching movements away from the mirroring axis, where they remained until visual feedback was removed. Participants never became able to compensate for the perturbation. This pattern of behavior is consistent with implicit adaptation occurring through direct policy updating but is difficult to reconcile with forward-model-based learning.

### Allowing participants to use strategy does not prevent error amplification to targets on the mirroring axis

It has been well established that learning to compensate for an imposed perturbation is not solely due to implicit adaptation, but is also attributable in part to explicit re-aiming of movements (Benson et al., 2011; Morehead et al., 2015; Taylor et al., 2014). To assess the possible effect of explicit re-aiming strategies, we ran a second group of participants which had an identical training schedule with one major difference: upon briefing participants about the nature of the upcoming mirror perturbation after the third block, we instructed them to do their best to bring the cursor through the target (rather than to ignore the cursor and bring their hand to the target, as for the first group). We refer to this group as the Free-aim group, in contrast to the first group, which we refer to as the Target-aim group. Similar to the Target-aim group, the Free-aim group showed an increase in errors for the on-axis targets, as shown in Figure 3D,E (2.7±0.2° in baseline vs. 18.0±2.1° in the last two blocks of adaptation, p <10^−5^). Despite the contrasting instructions, behavior was not significantly different between the two groups (2-way ANOVA for subject-averaged absolute errors, using time (baseline vs. late learning) and group (Free-aim vs. Target-aim) as factors; main effect of time (p<10^−8^) but no effect of group (p = 0.86), nor a time x group interaction, p = 1.00).

Prior to the 8^th^ block, participants were instructed to disengage any deliberate compensation strategies, and instead aim their hand through the cursor, and visual feedback was removed for this block. Participants exhibited aftereffects (absolute reaching angle on the first post-learning cycle: 15.3±1.8° vs. 4.3±0.4° for the no-visual-feedback baseline (block 2), p <10^−4^) and the magnitude of the aftereffects matched the amount of late adaptation, as suggested by a linear relationship with a slope close to 1 (red in Figure 3B, linear fit between last adaptation block and the first aftereffect cycle: slope = 0.87 (95% c.i.: [0.67, 1.07]), R^2^ = 0.90, p < 10^−5^). To systematically compare aftereffects in the two groups, we performed a 2-way ANOVA for subject-averaged absolute errors, using time (last adaptation block vs. first aftereffect cycle) and group (Free-aim vs. Target-aim) as factors; there was no main effect of time (p=0.57), group (p = 0.74), or any time x group interaction (p = 0.87).

These aftereffects demonstrate that implicit adaptation occurred at the on-axis targets, and that this errant learning could not be countered by the explicit system during learning even though participants were allowed to adopt a strategy. This stands in contrast to the relative ease by which participants minimized error in the off-axis targets, as illustrated by smaller absolute errors than the on-axis targets (7.9±1.5° in the last two adaptation blocks for the off-axis targets, compared to 18.0±2.1° for the on-axis targets, p =0.0042), suggesting that the effective use of explicit strategies was limited to the off-axis targets.

### Implicit adaptation drives movement away from mirroring axis at off-axis targets

The results from Experiment 1 suggested that implicit adaptation was unable to compensate for the imposed mirror reversal. However, a limitation of Experiment 1 is that, given their baseline biases to one side of other of the mirroring axis (Figure 3C), participants might never have sampled movements that would have led to successful target acquisition, and thus would not have been able to learn an accurate forward model prediction for those movements and therefore could not have known to select them. It is possible, therefore that forward-model-based learning might have similarly failed under the protocol of Experiment 1.

To address this limitation, we ran a second experiment, Experiment 2, which employed 12 different target directions equally spaced around the circle, instead of the four target directions used in Experiment 1. Targets were symmetrically positioned across the mirroring axis (Figure 4A) to ensure that participants gained experience making movements that would serve as the appropriate solution for every target. Participants should therefore have been able to learn an accurate forward model, and forward-model-based learning ought to have been able to successfully compensate for the mirror reversal.

**Figure 4.**
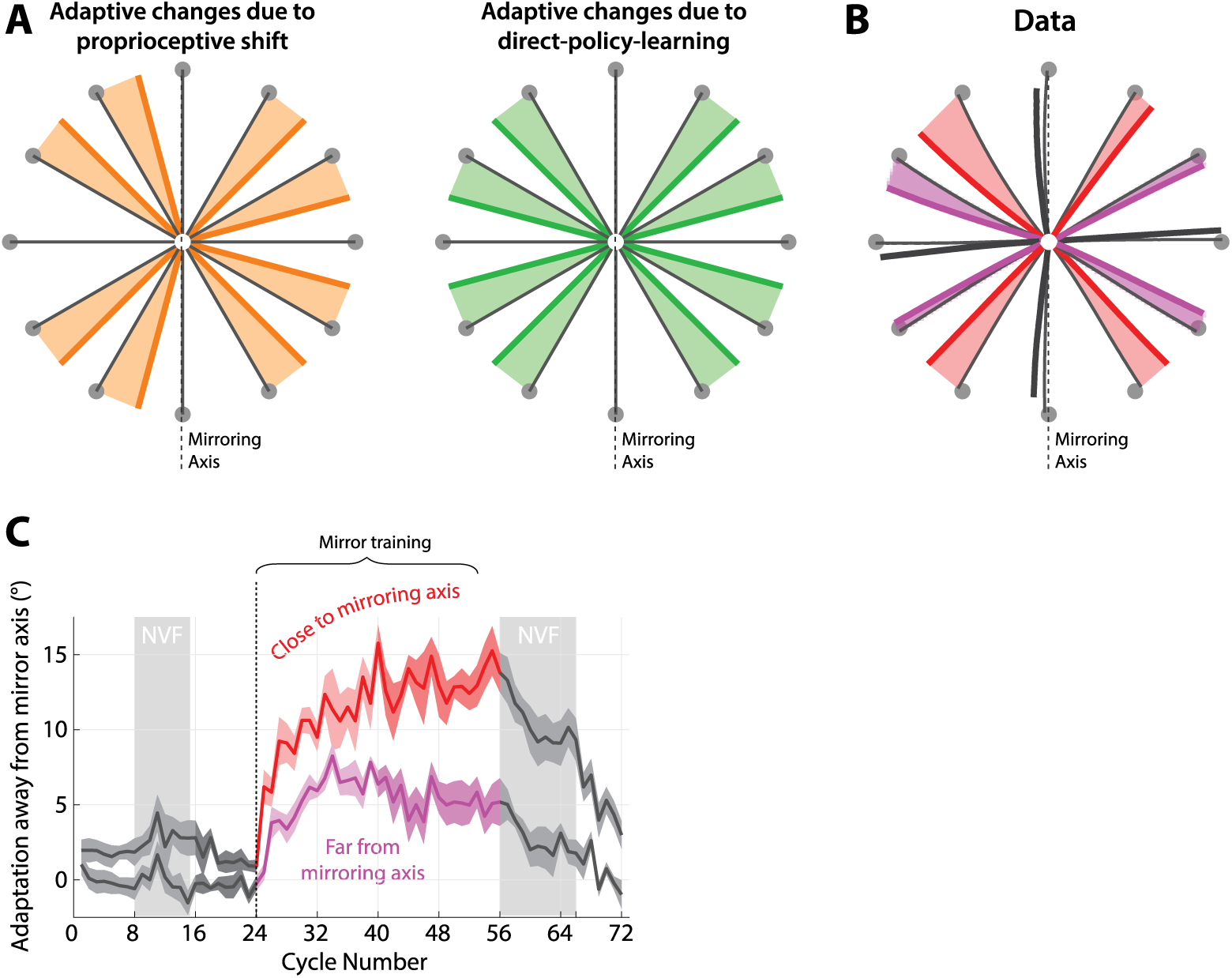
Adaptive changes to a mirror reversal when reaching to off-axis targets are consistent with direct policy learning, not forward-model-based learning or proprioceptive shifts. **A:** Expected pattern of changes in reach direction based on proprioceptive shift of the starting position (thick orange lines) vs. direct policy learning adaptive changes (thick green lines). If the observed adaptive changes were due to a leftward shift in the perceived location of the hand at the start of each movement (Sober & Sabes, 2003) would lead to systematic shift of all movements towards the rightward target. By contrast, drifts due to errant direct policy learning would always be directed away from the mirroring axis. Shading covers the area between baseline and adapted trajectories, to highlight the predicted direction of changes. **B:** Average reaching trajectories in Experiment 2 during baseline (thin black lines) and asymptote training (last two blocks; thick lines). Red: off-axis targets close to the mirroring axis; magenta: off-axis targets far from the mirroring axis; gray: targets on the mirroring axis or perpendicular to it. Arrows show the direction of adaptive changes as a result of training. Shading highlights these changes in a similar way to (A). **C:** Trial-to-trial adaptation away from the mirroring axis for targets close to and far from the mirroring axis (red and magenta, respectively, as in A; darker shades illustrate the last block of training). Baseline adaptation levels are highlighted in dark gray; shading represents SEM. NVF indicates no visual feedback.

This experiment also addresses another potential confound in Experiment 1: the pattern of directional biases we observed could have been the result of a shift in the proprioceptive estimate of the starting position of the hand (Sober & Sabes, 2003), rather than implicit adaptation. For example, if the hand is perceived to be to the left of the starting position, this would lead to a clockwise bias at the “up” target and a counter-clockwise bias at the “down” target, as illustrated in Figure 4C, which we indeed observed for many of our subjects, as in the example given in Figure 2C. Other patterns of shifts might also be explicable in terms of misestimated initial limb posture (Sober & Sabes, 2003). This potential alternative explanation for the results of Experiment 1 could be ruled out by assessing behavior across a broader range of targets (Figure 4A,B).

Participants completed 32 cycles of exposure to the mirror reversal with this set of targets. We focused our analysis on targets along the eight non-cardinal directions. At these targets, under forward-model-based learning, participants should eventually be able to learn appropriate compensation, whereas under direct policy learning they should not (Figure 4A).

We found that, for targets on the eight non-cardinal directions, there was a clear change in reach direction away from the mirroring axis (targets 30° away from mirroring axis, red in Figure 4BC: 1.4±0.4° vs. 13.1±1.1° for baseline vs. asymptote adaptation, p = 0.000002; targets 60° away from mirroring axis, magenta in Figure 4BC: −0.4±0.3° vs. 5.3±0.9°, p = 0.0001), and participants never became able to compensate for the perturbation – in line with predictions of direct policy learning but inconsistent with forward-model-based learning. Our Experiment 2 data also cannot be explained by a proprioceptive shift, since movements became biased away from the mirroring axis in both the left and right half-plane, inconsistent with a proprioceptive shift which would predict shifts in a similar direction for targets on opposite sides of the mirroring axis (Figure 4A).

### Unstable learning cannot be explained by limited extent of forward-model based learning

An important and well-documented characteristic of implicit adaptation is the fact that it is limited in its extent: under a visuomotor rotation, implicit adaptation saturates at about 15-25 degrees (Bond & Taylor, 2015; Kim et al., 2018; Morehead et al., 2017). Our off-axis targets in Experiment 2, are 30 or 60 degrees away from the mirroring axis, meaning that a forward model would need to change its output by 60 or 120 degrees, respectively. This is far beyond the established limits of implicit adaptation.

The issue of saturation of learning raises an important alternative explanation for the results of Experiment 2. We have assumed that, under forward-model-based learning, participants should be able to eventually learn an accurate forward model under a mirror-reversal and should therefore eventually be able to apply appropriate compensation. At intermediate points during learning, however, a partially adapted forward model would actually lead participants to select actions further away from the correct solution than at baseline – similar to our expectations for direct policy updating (Figure 5A). Saturation of learning might prevent participants from progressing beyond this intermediate state, potentially accounting for participants’ failure to compensate for the mirror-reversal and the apparent instability of learning.

**Figure 5:**
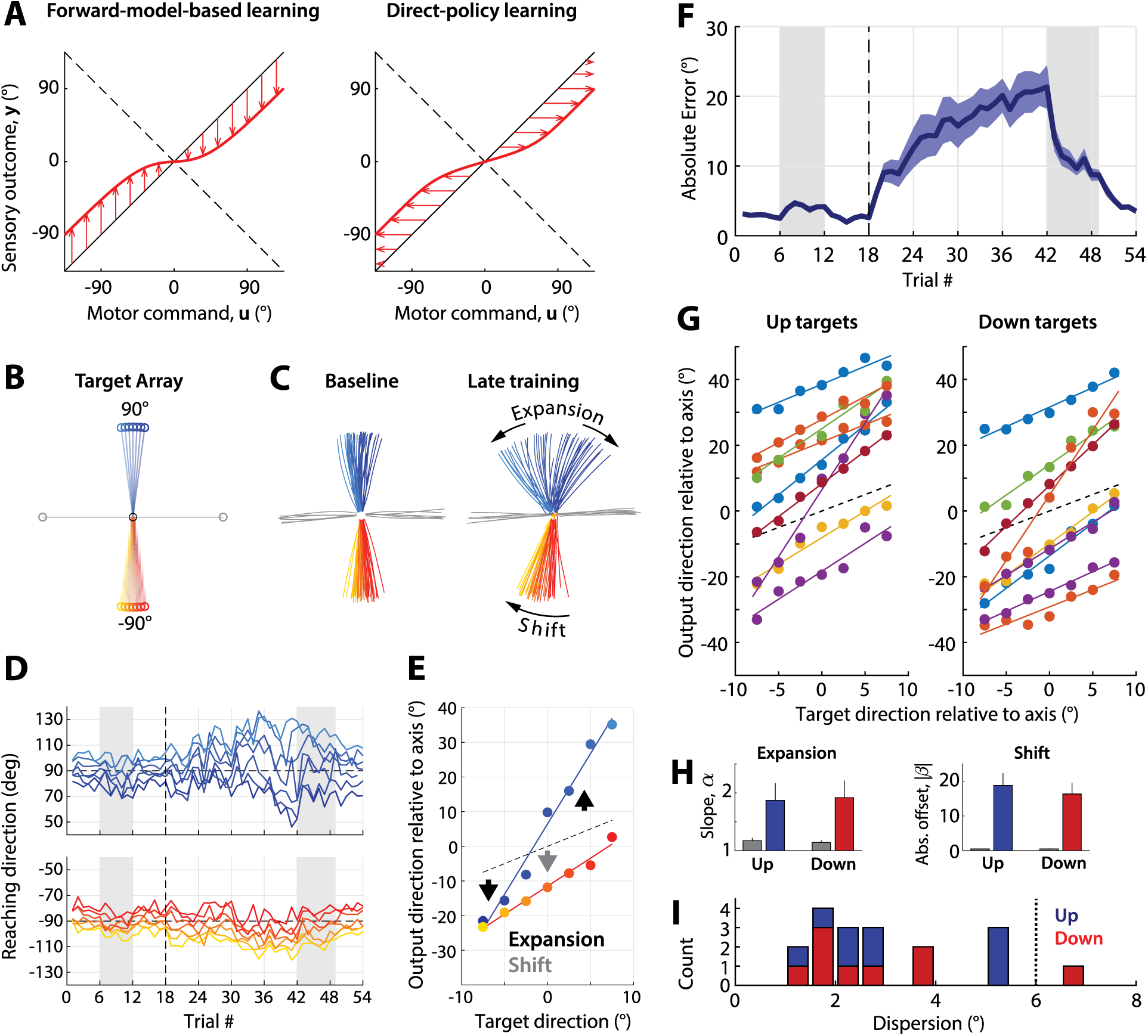
Instability of learning for a narrow range of targets. **A: Left:** Illustration of unstable learning predicted by forward-model-based learning. Learning occurs in the appropriate direction (arrows point towards the dashed line representing the new mapping) but a combination of saturation and broad generalization could give rise to a forward model that closely resembles the inverse of a policy arrived at by unstable direct policy learning (**right**). **B:** Target array used in Experiment 3. Targets are shown connected to the starting position (black) for clarity. **C:** Trajectories from an example participant during baseline (left) and the last training block (right). **D:** Trial-to-trial adaptive changes for the same participant (blue shades: “up” targets; red shades: “down” targets). **E:** Relationship between target direction and direction of motor output during baseline (empty circles) vs. late learning (filled circles) for the participant in B. The “up” targets predominantly show expansion whereas the “down” targets predominantly show a shift. **F:** Absolute error as a function of trial averaged across all participants. Shading indicates SEM. **G:** Relationship between target direction and direction of motor output during late learning for all participants, indicating a strong linear relationship. **H:** Estimated expansion and shift from baseline (gray) to asymptote adaptation, based on the linear relationship between target and motor output direction. Left: slope, increase of which (>1) indicates expansion; Right: (absolute) offset, which indicates a shift. **I:** Histogram of dispersion of differences in reach direction across neighboring targets for “up” (blue) and “down” (red) targets, indicating the extent of nonlinearity of the relationship between target location and motor output.

To examine this possibility, we ran a further experiment, Experiment 3, which focused on targets in a narrow but densely sampled area around the mirroring axis (between −7.5° and 7.5° about the 90 and 270 directions, in 2.5° increments, see Figure 5B). Potential limitations in adapting the forward model of around 15-25° should not preclude accurate learning of the forward model within this narrow range of targets, since the largest adaptive change required is just 15° – in contrast to Experiment 2 where the adaptive change required was at minimum 60° – far in excess of the apparent limits of implicit adaptation. Forward-model-based learning thus ought to be able to successfully adapt within this narrow range of targets, even given possible limited extent of learning. In contrast, we expected instability under direct policy learning.

Participants’ behavior was still unstable for this set of targets, with absolute error increasing as training progressed (2.6±0.1° during baseline vs. 19.3±2.6° during the last training block, p=0.0002, Figure 5F), consistent with our predictions for direct policy learning. Contrary to our expectations, we observed two distinct regimes of instability in participants’ behavior: in some instances, trajectories to opposing targets diverged away from the mirroring axis (*expansion*, Figure 5C-E), as we expected. In other instances, however, trajectories to targets on both sides of the mirroring axis moved roughly in parallel towards one direction or the other (*shift*, Figure 5C-E). Some participants exhibited mixtures of these phenomena, i.e. a simultaneous shift and expansion, and patterns of behavior for upward and downward sets of targets for the same participant were also not necessarily the same.

Regardless of the specific pattern of divergence, we found that there was always a strong the relationship between target and motor output (Figure 5G; R^2^ values for linear fit all >0.87). Linear fits allowed us to quantify expansion as an increase in the slope of this relationship, and shift as an offset (Figure 5H). Across all participants we found both a significant slope increase (upward targets: 1.17±0.05 in baseline vs. 1.87±0.30 in asymptote, p = 0.032; downward targets: 1.14±0.04 in baseline vs. 1.92±0.30 in asymptote, p = 0.029) and increase in the absolute offset (“up” targets: 0.5±0.1° in baseline vs. 18.8±3.5° in asymptote, p = 0.0010; “down” targets: 0.5±0.1° in baseline vs. 16.3±3.2° in asymptote, p = 0.0013) as illustrated in Figure 5H.

Although the instability of learning across even a narrow range of targets appears inconsistent with forward-model-based learning, we reasoned that it might be possible that interference (generalization) of learning either side of the mirroring axis might have impaired forward-model-based learning: clockwise learning at a target on one side of the axis might be offset by generalization of counterclockwise learning at a target on the opposite side. This interference might prevent participants from ever being able to learn an accurate forward model. To examine whether such an explanation could credibly account for the behavior we observed in Experiment 3, we simulated behavior in the protocol tested in Experiment 3 using either forward-model-based or direct policy learning. Our simulations were based on standard state-space models of learning employing a linear combination of Gaussian basis function (Donchin et al., 2003), with saturation implemented by limiting the weights associated with each basis function (See Materials and Methods for details).

We observed both expansions and shifts in simulations of both models. The extent and likelihood of expanding or shifting depended on the exact sequence of targets experienced as well as on the model parameters (learning rate ***η***, basis function width ***σ***, and extent of saturation ***u***_***asymptote***_; see Figure 6 for examples). However, only the simulations based on direct policy learning predicted the preserved linear relationship we observed between target direction and movement direction (Figure 6C,D versus G,H). Simulations assuming forward-model-based learning predicted non-linear patterns of behavior with often abrupt differences in motor output for neighboring targets (e.g. Figure 6C, left). These abrupt transitions are a direct consequence of using the forward model to select a motor command given the target – particularly when the forward model predicts multiple potential actions yielding the same outcome.

**Figure 6:**
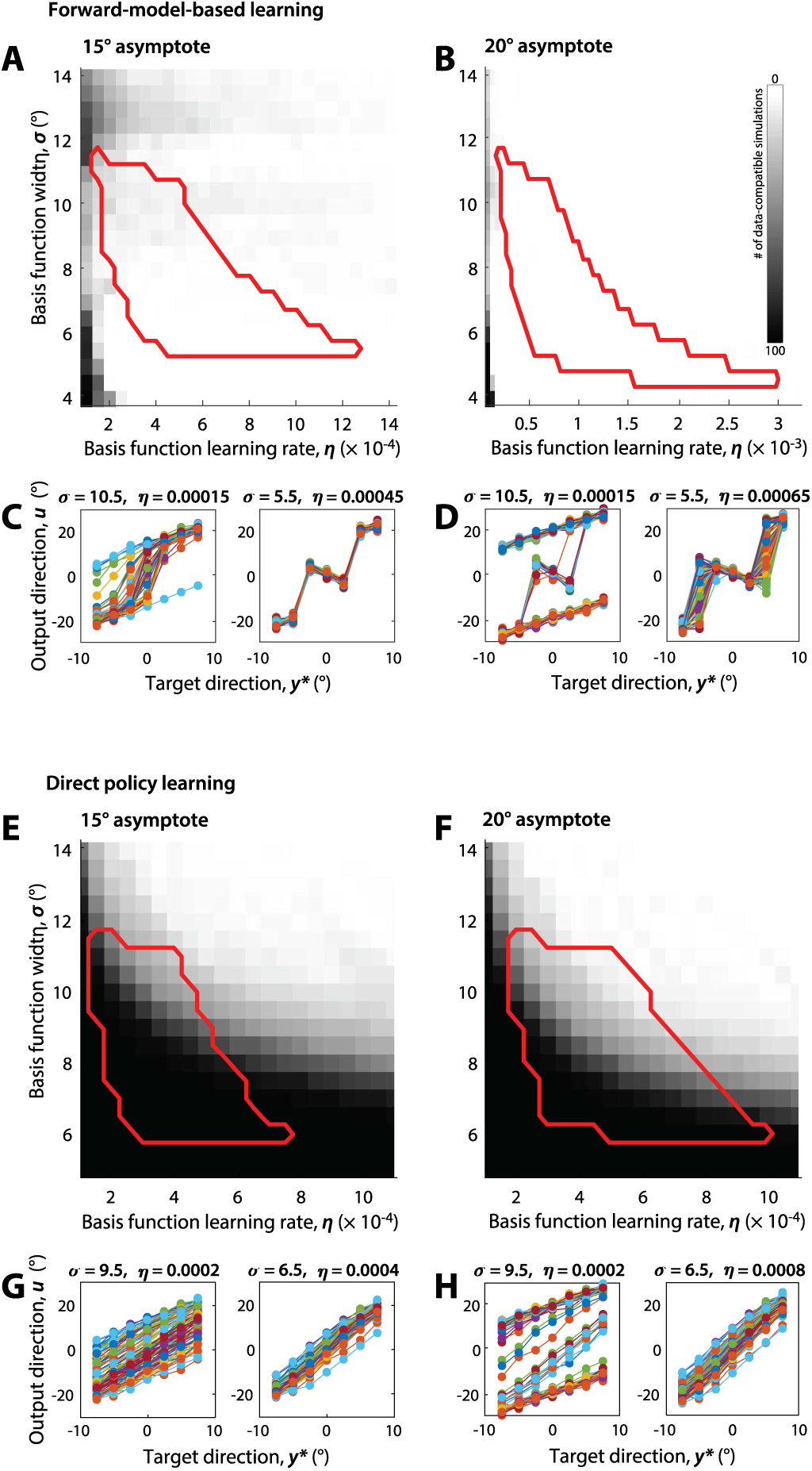
Simulations of forward-model-based learning and direct policy learning. **A-D:** Simulations of forward-model-based learning are inconsistent with the data and well-characterized attributes of implicit adaptation. The colormap indicates the # of pseudorandom simulations (out of 100) that resulted in expansion (slope >1.25) and smooth behavior (dispersion of differences across targets <6, see Figure 5H) as a function of learning function width and rate. The red border encloses parameter values consistent with learning and generalization of visuomotor adaptation based on (Morehead et al. 2017) (**A**: simulations using a 15 degree asymptote; **B**: simulations using a 20 degree asymptote). Panels **C** and **D** show simulated target vs output relationships after learning approaches a steady state (cycles 13-24, to match Experiment 3), for different parameter values (all 100 simulations for each parameter combination are shown in each panel). Note three distinct modes of behavior: shift (moving the target/output relationship one direction or the other without appreciably changing the slope); learning the inverted map, at least close to the mirroring axis (negative slope around 0, as in C, right); and in some cases expansion for all targets, (e.g. in some simulations in C, left); however, that expansion is defined by a characteristic S-shape in the target/output relationship, a form incompatible with our data (compare to Figure 5H). **E-H**: same as **A-D** but for direct policy learning, illustrating a wide array of model parameters compatible with our data.

To more systematically determine whether either class of model could account for our experimental data, we systematically varied the parameters of the simulated models (learning rate ***η***, basis function width ***σ***, and extent of saturation ***u***_***asymptote***_) over a wide range and assessed whether each model could qualitatively reproduce two of the main features of our data shown in Figure 5G,H: expansion, characterized by an increase in the slope of the relationship between target and asymptotic motor output direction, and a strong linearity in the same relationship. To assess whether simulation results reflected the linearity and smoothness of this relationship as we found in the data, we characterized the simulated target/output relationships in terms of the dispersion (standard deviation) of differences between neighboring targets, ***D***. This metric better captured the systematic nonlinearities in behavior predicted by forward-model-based learning than simply computing squared residual error. We considered a ***D*** above 6 to be inconsistent with our data (based on distribution observed in the data, Figure 5I). Furthermore, we assessed whether the model predictions were consistent with well-established properties of implicit adaptation, specifically the rate of adaptation, maximum extent of adaptation, and generalization to neighboring directions when implicit adaptation is driven by a constant error under an error-clamp paradigm (Morehead et al., 2017).

For forward-model-based learning, there was no overlap between parameters that could qualitatively account for our data under mirror-reversal, and parameters that could account for error-clamp behavior; simulation runs that predicted a linear expansion of motor output were very rare for an imposed asymptote of 15° (Figure 6A) and non-existent for asymptotes of 20° (Figure 6B) and 25° (Supplementary Figure 4). By contrast, there was broad overlap for the direct-policy-learning model between parameters that accounted for our data and parameters that accounted for error-clamp behavior (Figure 6E,F). We thus conclude that forward-model-based learning cannot account for our experimental observations, while they are naturally accounted for by the direct-policy-updating model.

## Discussion

Two potential mechanisms have previously been proposed for error-driven motor adaptation. One posits that observed errors are used to update a forward model which predicts the outcome of efferent motor commands, and that this forward model is then used to guide selection of future actions (forward-model-based learning). An alternative theory is that observed errors are used to directly update a policy that specifies motor output (i.e. a reach direction) for each possible target location (direct policy learning). Which of these theories best describes the underlying mechanism of adaptation has never been adequately resolved. Forward-model-based learning has become a widely accepted theory of adaptation. However, evidence for this is largely circumstantial – we know that forward models exist and can be altered through experience, and there is strong evidence that adaptation of motor output is driven by sensory-prediction errors, which is precisely the error signal one would expect to drive updating of a forward model.

Our experiments sought to identify the mechanism of adaptation by carefully examining behavior. We focused on the phenomenon of implicit adaptation under mirror-reversal – a phenomenon rarely considered in discussions of mirror reversal, since the mirror paradigm it is overwhelmingly associated with other learning mechanisms (Gutierrez-Garralda et al., 2013; Telgen et al., 2014), in spite of data clearly suggestive of implicit adaptation under mirror reversal (Lillicrap et al., 2013). This implicit adaptation under mirror-reversal that we isolate and observe bears the same hallmarks of implicit adaptation established in other paradigms, and therefore likely represents the same learning mechanism. Mirror reversal, however, serves as a critical condition under which forward-model-based learning can be distinguished from direct policy learning: whereas forward-model based learning ought to be able to adapt to a mirror reversal, direct policy learning will not. Though the results of our first experiment seemed inconsistent with forward-model-based learning, we considered numerous potential explanations as to how forward-model-based learning might have led to the observed behavior. We ran Experiment 2 to address the possibility that failure to compensate for the mirror-reversal in Experiment 1 was due to participants not having an opportunity to learn accurate forward model predictions for the relevant actions. We ran Experiment 3 to address the possibility that failure to compensate for the mirror-reversal in Experiment 2 was due to the limited extent to which a forward model could be recalibrated. Finally, we ran simulations to address the possibility that limited extent of learning, coupled with destructive interference between targets either side of the mirroring axis might have impaired the ability to compensate for the mirror-reversal under forward-model-based learning. Despite these efforts, we found no evidence to support the theory that implicit adaptation is driven by updating of an internal forward model which is then used to guide action selection. Instead, our findings are strongly consistent with the theory that adaptation occurs through direct updating of a control policy.

We would like to emphasize that the goal of our experiments was not to study how participants learn to compensate for a mirror reversal of visual feedback (which has been extensively examined elsewhere (Gritsenko & Kalaska, 2010; Lillicrap et al., 2013; Telgen et al., 2014; Wilterson & Taylor, 2019)). Instead, we used a mirror reversal to probe the properties of implicit adaptation. It has, however, been suggested that participants learn to compensate for mirror-reversal through a different mechanism than for visuomotor rotation (Telgen et al., 2014; Yang et al., 2020). Does this mean that the implicit learning we observed under mirror reversal might be qualitatively different than that which operates under a rotation perturbation? We consider this possibility very unlikely. Implicit adaptation seems to involuntarily adjust motor output in response to sensory prediction errors, and experienced sensory prediction error itself carries no information about the kind of perturbation that gave rise to it. To recruit distinct implicit adaptation systems for different types of perturbations, the motor system would need to somehow implicitly recognize the kind of perturbation that gave rise to the sensory prediction error. Furthermore, if the motor system were to use forward-model-based learning under a visuomotor rotation but switch to an alternate adaptation mechanism under a mirror reversal, this would be paradoxical since it would mean that the motor system switches away from a learning mechanism that could cope with mirror reversal (forward-model-based learning) in favor of one that cannot. Thus, we believe that the implicit adaptation we observed under a mirror-reversal is the same as is engaged under a visuomotor rotation. Ultimately, the qualitative differences between learning under a mirror-reversal and learning under a visuomotor rotation likely arise *because* implicit adaptation is incapable of learning to compensate for a mirror reversal and participants must therefore engage other learning mechanisms with qualitatively different properties.

### Relation to previous work on mirror-reversal learning

Our core observation – that adaptive responses exacerbate errors under a mirror reversal, rather than reducing them – has been demonstrated many times previously (Abdelghani & Tweed, 2010; Kasuga et al., 2015; Lillicrap et al., 2013). Similar to our interpretation, these previous studies explained this mal-adaptation in terms of task errors driving updates to a controller in a fixed manner (Abdelghani et al., 2008; Abdelghani & Tweed, 2010; Lillicrap et al., 2013). This work did not, however, distinguish between implicit and explicit contributions to learning. In principle, mal-adaptation could be attributable to misguided explicit compensation, rather than implicit adaptation. Our interpretation that the increase in errors is attributable to implicit adaptation is, however, recapitulated by a recent study examining implicit and explicit contributions to learning under mirror-reversal by collecting aiming reports (Wilterson & Taylor, 2019) (though the veracity of such aiming reports for assessing implicit learning has recently been called into question (Maresch & Donchin, 2019)). Several other studies have also shown that, although error tends to increase during the first few trials of exposure to a mirror reversal, errors start to decrease later in learning (Lillicrap et al., 2013; Wilterson & Taylor, 2019). This reversal may be attributable to learning a revised sensitivity derivative linking observed error to a corrective update (Abdelghani & Tweed, 2010; Kasuga et al., 2015) or to downregulation of implicit adaptation in favor of an explicit solution. The results of Experiment 1a suggest, however, that implementing an explicit strategy along the mirroring axis might be difficult – perhaps because explicit compensation also tends to be applied in an inappropriate direction following principles similar to the direct policy learning model we consider for implicit adaptation.

### Forward-model-based learning

Forward-model-based learning was originally proposed by Jordan and Rumelhart as a simple and effective solution to the distal error problem (Jordan & Rumelhart, 1992). The distal error problem is a major disadvantage of direct policy learning: errors are observed in task coordinates but in order to update the controller it is necessary to know the error in the outgoing motor command. Observed errors must therefore be translated from task space to motor space before they can be used to update the controller. This translation requires precise knowledge of the relationship between motor commands and task outcomes (Wolpert et al., 2001), which is tantamount to knowing what actions to take in the first place. The key insight from Jordan and Rumelhart’s work was that, translation of errors to the motor command space is not necessary when learning a forward model. Thus a fruitful general strategy for learning a controller is to first focus on learning a forward model.

Although the benefits of forward-model-based learning have influenced thinking about human motor learning, the question of how exactly a learned forward model might influence action selection has been neglected. One possibility is that the forward model could be used to simulate the outcomes of different motor commands and then selecting the one with the best outcome (Haruno et al., 2001; R Christopher Miall & Wolpert, 1996; Wolpert & Kawato, 1998). Alternatively, a learned forward model might be used to appropriately translate observed task errors into errors in motor errors (Jordan & Rumelhart, 1992) to serve as a teaching signal for updating a policy. This latter approach has the advantage that the forward model need not be perfectly learned in order to successfully improve action selection. It is worth noting, however, that similar learning of “sensitivity derivatives” relating task errors and motor errors can be learned through experience without needing to learn a forward model at all (Abdelghani et al., 2008)).

While this latter formulation does not necessarily require that the forward model is learned perfectly, it does require that the forward model is learned before a suitable controller can be learned. In line with this idea, it has been argued that forward-model prediction during an adaptation task is acquired more rapidly than the controller (Flanagan et al., 2003) using grip-force / load-force coupling as the indicator of prediction. However, it has previously been shown that, in a similar adaptation task, early adjustments of grip force may be driven by components non-specific to the adaptation task, but which would give the appearance of rapid grip force adaptation (Hadjiosif & Smith, 2015). Thus, despite some claims, there is little evidence to suggest that a forward model is learned before learning a policy to perform the task.

Other than direct policy learning, an alternative architecture that would also have difficulty adapting to a perturbation characterized by a flipped sensitivity derivative, like a mirror reversal, is the *feedback-error-learning model* (Albert & Shadmehr, 2016; Kawato, 1990; Kawato & Gomi, 1992). The feedback-error-learning hypothesis posits that feedback motor commands act as a training signal for updating the controller (since the correction is a motor command that ought to be similar to the error in the initial motor output). This scheme would also be unable to adapt under a mirror reversal, since the feedback controller would also fail to account for the flipped sensitivity derivative; feedback motor commands and, consequently, changes in feedforward motor commands, would thus be in the opposite direction from what is needed. In our experiments, however, we used a task design in which feedback corrections were minimized (if they occurred at all), making feedback-error learning a very unlikely explanation for the adaptive changes we observed.

### Neural basis of implicit adaptation

A longstanding theory of the neural basis of adaptation suggests that internal models for motor control are encoded in the strengths of synaptic connections between cerebellar Purkinje cells and parallel fibers. When an error is experienced, strong discharges from climbing fibers (originating in the inferior olive) carry error information to the corresponding Purkinje cells, resulting in complex spike activity in the Purkinje cells. Complex spike activity may in turn alter the strength of parallel-fiber connections in such a way as to update the internal model (Albus, 1971; Marr & Thach, 1991). Recent findings about the role of cerebellar Purkinje cells in adaptation shed more light in how such a mechanism might work (Herzfeld et al., 2018). Specifically, it was found that the error-signaling complex spikes result in changes to simple spike activity in Purkinje cells only along the dimension of the particular cell’s preferred error, with cells responsive to leftward errors only resulting in rightward shifts in motor output and vice versa. These findings are not necessarily incompatible with direct policy learning, as such hard-wired associations between error direction and correction direction could reflect a fixed sensitivity derivative to errors.

### Is there any role for a forward model in adaptation?

While we have argued that implicit adaptation does not result from the inversion of an updated forward model, it is important to clarify that we are not arguing against the existence of forward models, or against the idea that they may play an important role in implicit adaptation. There is substantial evidence, both behavioral and neurophysiological, that the brain uses an internal forward model to maintain an estimate of the state of the body (Bhanpuri et al., 2013; Ebner & Pasalar, 2008; Herzfeld et al., 2018; R. Chris Miall et al., 2007; Wagner & Smith, 2008), and that these estimates can be updated in the presence of a perturbation (Cressman & Henriques, 2010; Izawa & Shadmehr, 2011; Synofzik et al., 2008). This state estimate is critical for maintaining stable control of the body in the presence of signaling delays.

Such forward models also appear to be important for learning. Implicit adaptation seems to be driven by sensory prediction error (Leow et al., 2018; Mazzoni & Krakauer, 2006; Taylor et al., 2014), which itself implies a sensory prediction which, presumably, arises as the output of a forward model. Therefore, even though changes to the forward model do not directly influence action selection, they may influence the way in which the policy is updated. This interdependence between forward model learning and policy learning could lead to interesting interactions. For instance, if updates to the controller are driven by sensory-prediction error, at some point sensory prediction error would reach zero, at which point there would no longer be any error signal to drive to changes in the controller. Such a scenario could potentially account for the apparent limited extent of adaptation, which saturates at about 20 degrees in the case of visuomotor perturbations (Bond & Taylor, 2015; Kim et al., 2018; Morehead et al., 2017; Taylor et al., 2014).

## Materials and Methods

### Participants and Ethics Statement

A total of 51 individuals were recruited for the study. Four participants were excluded prior to analysis: three because they didn’t complete the study due to time constraints, and one due to inability to follow instructions. This resulted in 47 participants (age: 23.3±5.2 (mean±standard deviation), 20 identifying as male, 26 identifying as female, 1 identifying as nonbinary), 12 in Experiments 1, 1a, and 3, and 11 in Experiment 2. Sample size was determined based on similar behavioral studies. Participants used their dominant arm for the task; four participants self-reported as left-handed; another was ambidextrous and used their right arm for the task. For left-handed participants, all data were flipped about the mirroring axis before further analysis. All participants had no known neurological disorders and provided informed consent before participating. Study procedures were approved by the Johns Hopkins University School of Medicine Institutional Review Board.

### Task details

Participants sat on a chair in front of a table, with their dominant arm resting on an air sled which enabled planar movement with minimal friction against the glass surface of the table. Targets (diameter: 10 mm) and a hand-controlled cursor (diameter: 5 mm) were presented on the plane of movement with the help of a mirrored display. Hand position was tracked at 130 Hz using a Flock of Birds magnetic tracking device (Ascension Technologies).

Participants performed shooting movements through targets positioned 12cm away from a central, starting position (Experiments 1 and 1a: four different targets arranged along the cardinal directions as illustrated in Figure 3B; Experiment 2: twelve different targets evenly spaced across the circle (every 30°, beginning at 0°), as illustrated in Figure 5A; Experiment 3: sixteen different targets, two horizontal ones (0° and 180° directions), seven around the 90° direction in 2.5° increments(82.5, 85, 87.5, 90, 92.5, 95, 97.5), and similarly seven around the 270° direction. When the cursor reached 12cm away from the starting position (equal to the target distance), its color would change and it would freeze for 0.5s to indicate how close the participant came to going through the target; afterwards, the participant was instructed to return to the starting position. During the return movement, we replaced cursor feedback with a circle, centered on the start location, whose diameter indicated the distance, but not exact location, of the (hidden) cursor, in order to avoid any learning during the return movement. To facilitate return movements, an air jet positioned above the start position blew a narrow stream of air downwards that participants could feel on their hand.

During the experiment, velocity was monitored by the experimenter. To ensure that participants were moving at a brisk speed, participants were encouraged to adjust their movement speed if they were being excessively slow or fast.

For all experiments, participants were informed of the nature of the mirror reversal right before the block in which it was first imposed. In Experiments 1, 2 and 3, participants were instructed to keep aiming their hand through the target (Target-aim groups) – even if that meant the cursor could deviate from the target. In Experiment 1a, participants were instructed to try to get the cursor through the target (Free-aim group).

All experiments consisted of 9 blocks, between which short rest breaks were given. The number of trials in each block depended on the experiment (Experiments 1 and 1a: 20 trials to each of the 4 targets; Experiment 2: 8 trials to each of the 12 targets; Experiment 3: 6 trials to each of the 16 targets). Depending on the block, one of three different types of visual feedback was used for the out movements: veridical cursor feedback (Blocks 1 and 3), visual feedback inverted about the y axis (mirror reversal, Blocks 4-7), and no visual feedback (Blocks 2 and 8), whereby cursor feedback was withheld in order to assess any baseline biases (in Block 2) or aftereffects of adaptation (in Block 8). Block 9 began without visual feedback, then transitioned to veridical visual feedback. In Experiments 1 and 1a, this transition happened after 5 cycles. For Experiment 2 and 3 participants in Experiment 3, due to an implementational error, this transition occurred in the middle of a cycle (after 20 trials), rather than after a complete cycle. However, this did not significantly affect the results since data from these trials were not part of our main analysis. For the remaining participants in Experiment 3 this was corrected so that the transition occurred after 16 trials (one cycle).

### Data analysis

Analysis was performed using Matlab (Mathworks, Natick MA). Position data were smoothed by filtering through a 3^rd^ order Savitzky-Golay filter with a window size of 9 samples (69ms). The main outcome variable extracted from the data was the reaching angle relative to the target direction, measured at target distance (12cm).

For Experiments 1 and 1a, we focused on adaptation for the two targets on the mirroring axis. For these targets, the direction of the (mal-adaptive) drift was inconsistent across participants. Therefore, to aggregate data across participants, we used absolute error as the primary measure of adaptive changes.

For Experiment 2, we focused on adaptation for the eight targets not in cardinal directions. These targets lied 30° or 60° away from the mirroring axis and used signed error as the primary measure of adaptive changes, flipping the sign as appropriate so that a positive error always indicated adaptive changes away from the mirroring axis.

For Experiment 3, we focused on the adaptive changes for the two sets of targets about the up/down directions. As in Experiment 1, we quantified learning in terms of the absolute error. However, to more precisely characterize the nature of adaptive changes we also fit a linear relationship between target direction and hand movement output. This yielded two parameters for each direction (upward/downward) for each participant: a slope and an offset. We refer to increases in the slope as *expansion*, and changes in the offset as a *shift*.

#### Data inclusion criteria

We analyzed trials along the horizontal targets to monitor adherence to the aiming instructions for Experiments 1, 2, and 3. For example, if the participant were aiming their hand through the target in spite of cursor errors as was the instruction in Experiment 1, the off-axis targets would show a cursor error of about 180°, whereas if they were not following the instruction the cursor error would be closer to 0°. This resulted in a bimodal distribution of reach directions along the horizontal targets, making it easy to detect trials in which participants failed to follow this instruction (See Supplementary Figure 1). We classified horizontal trials as out of line with instructions if the movement direction was closer to the opposing target (i.e. hand direction error >90°). Errors of this kind were rather rare (average on participants included in analysis: 2.4%), with the exception of four participants (one in Experiment 1, one in Experiment 2, and two in Experiment 3) who had such errors on more than 10% of horizontal-target trials during mirror adaptation. We excluded these four participants from our final analysis. A supplementary analysis including these participants is provided in Supplementary Figure 2 showing that they behaved in a way similar to the rest of the population, and thus retaining their data would not have altered our conclusions.

In addition, one participant in Experiment 3 was excluded due to erratic reaching behavior, which was in line with neither forward-model-based nor direct-policy-update-based learning but likely reflected large trunk postural adjustments during the experiment. This participant’s data are shown and discussed in Supplementary Figure 3.

Finally, for trials to targets other than the horizontal ones, in all experiments, we excluded as outliers trials for which the absolute error was >75°; these trials constituted a very small fraction of the total (0.46%).

#### Statistics

Within each group, we compared adaptation during baseline with adaptation during late learning using paired 2-tailed t-tests. To investigate the effect of instruction (Experiment 1 vs 1a) we used a 2×2 ANOVA with group (target-aim vs. free-aim group) and time (baseline vs. late learning) as factors, whereas we used a similar 2×2 ANOVA to investigate the relationship between aftereffects and late adaptation with group (target-aim vs. free-aim group) and time (last training block vs. first aftereffects cycle) as factors.

### Simulating direct-policy-update and forward-model based learning

#### Forward-model based learning

We assume that the forward model, 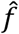, which approximates the relationship, *f*, between motor commands, *u* and their sensory outcome, *y* = *f*(*u*), is constructed as a linear sum of non-linear basis functions:

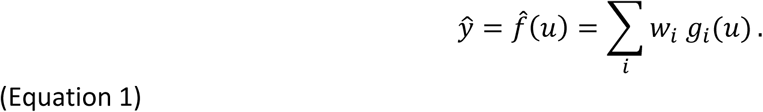

Here, individual basis functions *g*_*i*_ are combined according to weights *w*_*i*_ which can change during adaptation to build an improved approximation to the perturbed sensorimotor map.

In our simulations, we modeled adaptation of the forward model by assuming the motor system is using gradient descent to minimize a function C of the error, such as

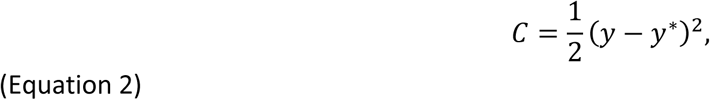

where y* is the desired sensory output. Assuming that the motor system tries to select a motor command that would yield the desired sensory outcome y* by inverting the forward model at *ŷ*, we can substitute (*ŷ* = *y*^∗^), which makes the learning rule that minimizes C equal to

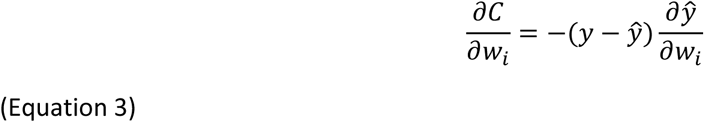

Or,

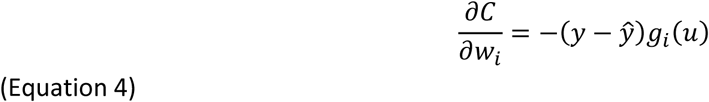

Thus, the learning update is given by the sensory prediction error, *y* − *ŷ*, multiplied by a learning rate *η*.

#### Direct-policy-update based learning

The control policy (inverse model) 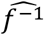 outputs the motor commands *u* to bring about a desired sensory change *y*^∗^, approximating the inverse of *f*:

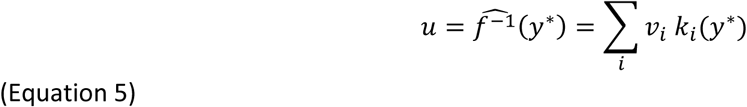

With the goal of minimizing error as in Equation 2, the learning rule that minimizes C becomes

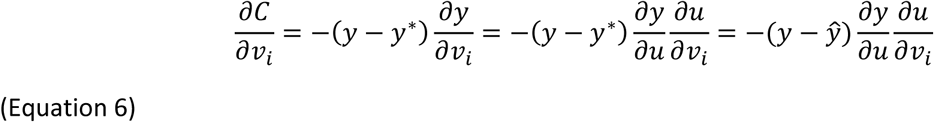

Here, 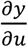 transforms the sensory error into a motor error, and is sometimes referred to as a *sensitivity derivative* (Abdelghani et al., 2008; Abdelghani & Tweed, 2010). Equation 6 can also be rewritten as

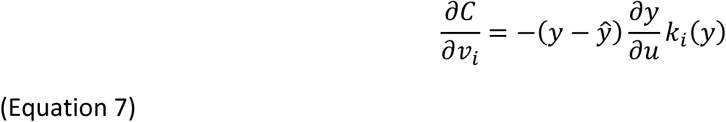

Thus, like in forward model learning, the direct-policy-update-based learning update will be countering he sensory prediction error, *y* − *ŷ*, times a learning rate *η*; however, this will also need to be multiplied by the sensitivity derivative 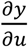. Without a mechanism to learn the sensitivity derivative, the motor system can only rely on simple rules about the sign and magnitude of the sensitivity derivative, i.e. here, 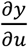 is a constant.

To simulate forward-model-based and direct policy learning, we modeled the learning basis functions as Gaussian kernels:

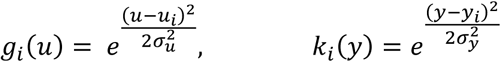

The centers of each basis function, *u*_*i*_ and *y*_*i*_ above, were uniformly distributed around the mirroring axis with a spacing of 0.05°. The widths of these kernels, *σ*_*u*_ and *σ*_*y*_ were left as open parameters.

To model saturation in learning, we saturated the basis function weights using a sigmoidal function:

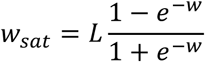

The asymptotic limit of adaptative changes in motor output, *u*_*out,asymptote*_ linearly scales with *L*:

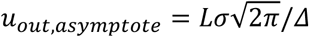

where *σ* is the width of the corresponding basis function (*g* or *k*, correspondingly) and *Δ* is the spacing between consecutive basis functions (0.05° in our simulations).

For forward-model-based learning we used sensory-prediction error to drive learning (*y* − *ŷ*); for direct-policy-update-based learning, we used task error (*y* − *y*^∗^).

To select an action in each trial, under forward-model-based learning, we selected the action which the forward model predicted would lead to the desired outcome. In cases where multiple motor commands were predicted to lead to the same expected sensory outcome, we chose the motor command closest to the origin.

To identify possible model parameters, we systematically examined behavior for a range of values for the *σ* and *η* parameters (Specifically, for forward-model-based learning: *σ* ∈ [4°, 14°] in 0.5° increments, and *η* ∈ [1,14] × 10^−4^ (for 15° asymptote simulations) or *η* ∈ [1,32] × 10^−4^ (for 20° asymptote simulations) in 0.5 × 10^−4^ increments; for direct policy learning: *σ* ∈ [5°, 14°] in 0.5° increments, and *η* ∈ [0.5,12] × 10^−4^ in 0.5 × 10^−4^ increments). For each pair of parameters (*σ, η*) and for each of the two models, we ran 100 simulations, each with a different random target order. Specifically, we randomized the order of the 7 targets within each cycle, for a total of 24 cycles (168 trials), to match the number of training cycles in Experiment 3. We also included motor noise in our simulations by adding zero-mean Gaussian noise to the outgoing motor commands, with a standard deviation of 3.2° based on baseline data. As in our analysis of Experiment 3, we used the last two “blocks” (last 12 cycles) as a measure of asymptote adaptation.

To classify which simulations led to expansion, we set a criterion that the resulting slopes of the target – output relationship should be greater than 1.25. To estimate values for (*σ, η*) that are compatible with the learning and generalization characteristics of visuomotor adaptation, we simulated the adaptation of both types of models to single-target adaptation to a 22.5° visual clamp (Morehead et al. 2017). We evaluated the width of the resulting generalization function, taking as compatible parameters the ones which led to a full-width-half-max between 40° and 80°, and speed of learning, taking as compatible parameters the ones which led to between 6 and 18 trials to reach 80% of asymptotic adaptation.

## Contributions

A.M. Hadjiosif and A.M. Haith conceived and designed the experiments; A.M. Hadjiosif performed experiments, analyzed the data and prepared all figures. A.M. Hadjiosif and A.M. Haith wrote the paper. All authors edited the paper.

## Acknowledgements

A.M. Hadjiosif is supported by the Sheikh Khalifa Stroke Institute.

## References

Abdelghani, M. N., Lillicrap, T. P., & Tweed, D. B. (2008). Sensitivity derivatives for flexible sensorimotor learning. Neural Computation, 20(8), 2085–2111.

Abdelghani, M. N., & Tweed, D. B. (2010). Learning course adjustments during arm movements with reversed sensitivity derivatives. BMC Neuroscience, 11(1), 150.

Albert, S. T., & Shadmehr, R. (2016). The neural feedback response to error as a teaching signal for the motor learning system. Journal of Neuroscience, 36(17), 4832–4845.

Albus, J. S. (1971). A theory of cerebellar function. Mathematical Biosciences, 10(1), 25–61. https://doi.org/10.1016/0025-5564(71)90051-4

Bastian, A. J. (2006). Learning to predict the future: The cerebellum adapts feedforward movement control. Current Opinion in Neurobiology, 16(6), 645–649.

Benson, B. L., Anguera, J. A., & Seidler, R. D. (2011). A spatial explicit strategy reduces error but interferes with sensorimotor adaptation. Journal of Neurophysiology, 105(6), 2843–2851.

Bhanpuri, N. H., Okamura, A. M., & Bastian, A. J. (2013). Predictive modeling by the cerebellum improves proprioception. Journal of Neuroscience, 33(36), 14301–14306.

Bhushan, N., & Shadmehr, R. (1999). Computational nature of human adaptive control during learning of reaching movements in force fields. Biological Cybernetics, 81(1), 39–60.

Bond, K. M., & Taylor, J. A. (2015). Flexible explicit but rigid implicit learning in a visuomotor adaptation task. Journal of Neurophysiology, 113(10), 3836–3849.

Cressman, E. K., & Henriques, D. Y. P. (2010). Reach Adaptation and Proprioceptive Recalibration Following Exposure to Misaligned Sensory Input. Journal of Neurophysiology, 103(4), 1888–1895. https://doi.org/10.1152/jn.01002.2009

Donchin, O., Francis, J. T., & Shadmehr, R. (2003). Quantifying Generalization from Trial-by-Trial Behavior of Adaptive Systems that Learn with Basis Functions: Theory and Experiments in Human Motor Control. Journal of Neuroscience, 23(27), 9032–9045.

Ebner, T. J., & Pasalar, S. (2008). Cerebellum predicts the future motor state. The Cerebellum, 7(4), 583–588.

Flanagan, J. R., Vetter, P., Johansson, R. S., & Wolpert, D. M. (2003). Prediction precedes control in motor learning. Current Biology, 13(2), 146–150.

Galea, J. M., Vazquez, A., Pasricha, N., Orban de Xivry, J.-J., & Celnik, P. (2010). Dissociating the roles of the cerebellum and motor cortex during adaptive learning: The motor cortex retains what the cerebellum learns. Cerebral Cortex, 21(8), 1761–1770.

Gritsenko, V., & Kalaska, J. F. (2010). Rapid online correction is selectively suppressed during movement with a visuomotor transformation. Journal of Neurophysiology, 104(6), 3084–3104.

Gutierrez-Garralda, J. M., Moreno-Briseño, P., Boll, M.-C., Morgado-Valle, C., Campos-Romo, A., Diaz, R., & Fernandez-Ruiz, J. (2013). The effect of Parkinson’s disease and Huntington’s disease on human visuomotor learning. European Journal of Neuroscience, 38(6), 2933–2940. https://doi.org/10.1111/ejn.12288

Hadjiosif, A. M., & Smith, M. A. (2015). Flexible Control of Safety Margins for Action Based on Environmental Variability. Journal of Neuroscience, 35(24), 9106–9121. https://doi.org/10.1523/JNEUROSCI.1883-14.2015

Haith, A. M., & Krakauer, J. W. (2013). Model-based and model-free mechanisms of human motor learning. In Progress in motor control (pp. 1–21). Springer, New York, NY. http://link.springer.com/10.1007/978-1-4614-5465-6_1

Haith, A., & Vijayakumar, S. (2009). Implications of different classes of sensorimotor disturbance for cerebellar-based motor learning models. Biological Cybernetics, 100(1), 81–95.

Haruno, M., Wolpert, D. M., & Kawato, M. (2001). Mosaic model for sensorimotor learning and control. Neural Computation, 13(10), 2201–2220.

Herzfeld, D. J., Kojima, Y., Soetedjo, R., & Shadmehr, R. (2018). Encoding of error and learning to correct that error by the Purkinje cells of the cerebellum. Nature Neuroscience, 21(5), 736.

Izawa, J., & Shadmehr, R. (2011). Learning from sensory and reward prediction errors during motor adaptation. PLoS Computational Biology, 7(3), e1002012.

Jayaram, G., Tang, B., Pallegadda, R., Vasudevan, E. V., Celnik, P., & Bastian, A. (2012). Modulating locomotor adaptation with cerebellar stimulation. Journal of Neurophysiology, 107(11), 2950–2957.

Jordan, M. I., & Rumelhart, D. E. (1992). Forward models: Supervised learning with a distal teacher. Cognitive Science, 16(3), 307–354.

Kasuga, S., Kurata, M., Liu, M., & Ushiba, J. (2015). Alteration of a motor learning rule under mirror-reversal transformation does not depend on the amplitude of visual error. Neuroscience Research, 94, 62–69.

Kawato, M. (1990). Feedback-error-learning neural network for supervised motor learning. In Advanced neural computers (pp. 365–372). Elsevier.

Kawato, M., & Gomi, H. (1992). A computational model of four regions of the cerebellum based on feedback-error learning. Biological Cybernetics, 68(2), 95–103.

Kim, H. E., Morehead, J. R., Parvin, D. E., Moazzezi, R., & Ivry, R. B. (2018). Invariant errors reveal limitations in motor correction rather than constraints on error sensitivity. Communications Biology, 1(1), 19. https://doi.org/10.1038/s42003-018-0021-y

Kitago, T., Ryan, S. L., Mazzoni, P., Krakauer, J. W., & Haith, A. M. (2013). Unlearning versus savings in visuomotor adaptation: Comparing effects of washout, passage of time, and removal of errors on motor memory. Frontiers in Human Neuroscience, 7. https://www.ncbi.nlm.nih.gov/pmc/articles/PMC3711055/

Kojima, Y., Iwamoto, Y., & Yoshida, K. (2004). Memory of Learning Facilitates Saccadic Adaptation in the Monkey. Journal of Neuroscience, 24(34), 7531–7539. https://doi.org/10.1523/JNEUROSCI.1741-04.2004

Krakauer, J. W., Ghez, C., & Ghilardi, M. F. (2005). Adaptation to visuomotor transformations: Consolidation, interference, and forgetting. Journal of Neuroscience, 25(2), 473–478.

Krakauer, J. W., Hadjiosif, A. M., Xu, J., Wong, A. L., & Haith, A. M. (2019). Motor learning. Comprehensive Physiology, 9, 613–663.

Krakauer, J. W., & Mazzoni, P. (2011). Human sensorimotor learning: Adaptation, skill, and beyond. Current Opinion in Neurobiology, 21(4), 636–644.

Krakauer, J. W., Pine, Z. M., Ghilardi, M.-F., & Ghez, C. (2000). Learning of visuomotor transformations for vectorial planning of reaching trajectories. Journal of Neuroscience, 20(23), 8916–8924.

Krakauer, J. W., & Shadmehr, R. (2006). Consolidation of motor memory. Trends in Neurosciences, 29(1), 58–64.

Leow, L., Marinovic, W., de Rugy, A., & Carroll, T. J. (2018). Task errors contribute to implicit aftereffects in sensorimotor adaptation. European Journal of Neuroscience, 48(11), 3397–3409.

Lillicrap, T. P., Moreno-Briseño, P., Diaz, R., Tweed, D. B., Troje, N. F., & Fernandez-Ruiz, J. (2013). Adapting to inversion of the visual field: A new twist on an old problem. Experimental Brain Research, 228(3), 327–339.

Malone, L. A., Vasudevan, E. V. L., & Bastian, A. J. (2011). Motor Adaptation Training for Faster Relearning. Journal of Neuroscience, 31(42), 15136–15143. https://doi.org/10.1523/JNEUROSCI.1367-11.2011

Maresch, J., & Donchin, O. (2019). Reporting affects explicit knowledge in visuomotor rotations in ways not measured by reporting. BioRxiv, 702290.

Marr, D., & Thach, W. T. (1991). A Theory of Cerebellar Cortex. In From the Retina to the Neocortex (pp. 11–50). Birkhäuser Boston. https://doi.org/10.1007/978-1-4684-6775-8_3

Mazzoni, P., & Krakauer, J. W. (2006). An implicit plan overrides an explicit strategy during visuomotor adaptation. Journal of Neuroscience, 26(14), 3642–3645.

McDougle, S. D., Ivry, R. B., & Taylor, J. A. (2016). Taking aim at the cognitive side of learning in sensorimotor adaptation tasks. Trends in Cognitive Sciences, 20(7), 535–544.

McNamee, D., & Wolpert, D. M. (2019). Internal models in biological control. Annual Review of Control, Robotics, and Autonomous Systems, 2, 339–364.

Miall, R. Chris, Christensen, L. O. D., Cain, O., & Stanley, J. (2007). Disruption of State Estimation in the Human Lateral Cerebellum. PLOS Biology, 5(11), e316. https://doi.org/10.1371/journal.pbio.0050316

Miall, R Christopher, & Wolpert, D. M. (1996). Forward models for physiological motor control. Neural Networks, 9(8), 1265–1279.

Morehead, J. R., Qasim, S. E., Crossley, M. J., & Ivry, R. (2015). Savings upon re-aiming in visuomotor adaptation. Journal of Neuroscience, 35(42), 14386–14396.

Morehead, J. R., Taylor, J. A., Parvin, D. E., & Ivry, R. B. (2017). Characteristics of Implicit Sensorimotor Adaptation Revealed by Task-irrelevant Clamped Feedback. Journal of Cognitive Neuroscience. http://www.mitpressjournals.org/doi/abs/10.1162/jocn_a_01108

Parrell, B., Agnew, Z., Nagarajan, S., Houde, J., & Ivry, R. B. (2017). Impaired feedforward control and enhanced feedback control of speech in patients with cerebellar degeneration. Journal of Neuroscience, 37(38), 9249–9258.

Porrill, J., Dean, P., & Stone, J. V. (2004). Recurrent cerebellar architecture solves the motor-error problem. Proceedings of the Royal Society of London. Series B: Biological Sciences, 271(1541), 789–796.

Shadmehr, R., & Mussa-Ivaldi, F. A. (1994). Adaptive representation of dynamics during learning of a motor task. Journal of Neuroscience, 14(5), 3208–3224.

Shadmehr, R., Smith, M. A., & Krakauer, J. W. (2010). Error correction, sensory prediction, and adaptation in motor control. Annual Review of Neuroscience, 33, 89–108.

Smith, M. A., Ghazizadeh, A., & Shadmehr, R. (2006). Interacting adaptive processes with different timescales underlie short-term motor learning. PLoS Biology, 4(6), e179.

Sober, S. J., & Sabes, P. N. (2003). Multisensory integration during motor planning. Journal of Neuroscience, 23(18), 6982–6992.

Synofzik, M., Vosgerau, G., & Newen, A. (2008). Beyond the comparator model: A multifactorial two-step account of agency. Consciousness and Cognition, 17(1), 219–239.

Taylor, J. A., Krakauer, J. W., & Ivry, R. B. (2014). Explicit and implicit contributions to learning in a sensorimotor adaptation task. Journal of Neuroscience, 34(8), 3023–3032.

Telgen, S., Parvin, D., & Diedrichsen, J. (2014). Mirror reversal and visual rotation are learned and consolidated via separate mechanisms: Recalibrating or learning de novo? Journal of Neuroscience, 34(41), 13768–13779.

Wagner, M. J., & Smith, M. A. (2008). Shared internal models for feedforward and feedback control. Journal of Neuroscience, 28(42), 10663–10673.

Wilterson, S. A., & Taylor, J. A. (2019). Implicit visuomotor adaptation remains limited after several days of training. BioRxiv, 711598.

Wolpert, D. M., Ghahramani, Z., & Flanagan, J. R. (2001). Perspectives and problems in motor learning. Trends in Cognitive Sciences, 5(11), 487–494.

Wolpert, D. M., & Kawato, M. (1998). Multiple paired forward and inverse models for motor control. Neural Networks, 11(7), 1317–1329.

Wong, A. L., & Shelhamer, M. (2011). Saccade adaptation improves in response to a gradually introduced stimulus perturbation. Neuroscience Letters, 500(3), 207–211. https://doi.org/10.1016/j.neulet.2011.06.039

Yang, C. S., Cowan, N. J., & Haith, A. M. (2020). De novo learning versus adaptation of continuous control in a manual tracking task. BioRxiv.

