## Supplementary Materials for "Did we get sensorimotor adaptation wrong? Implicit adaptation as direct policy updating rather than forward-model-based learning"

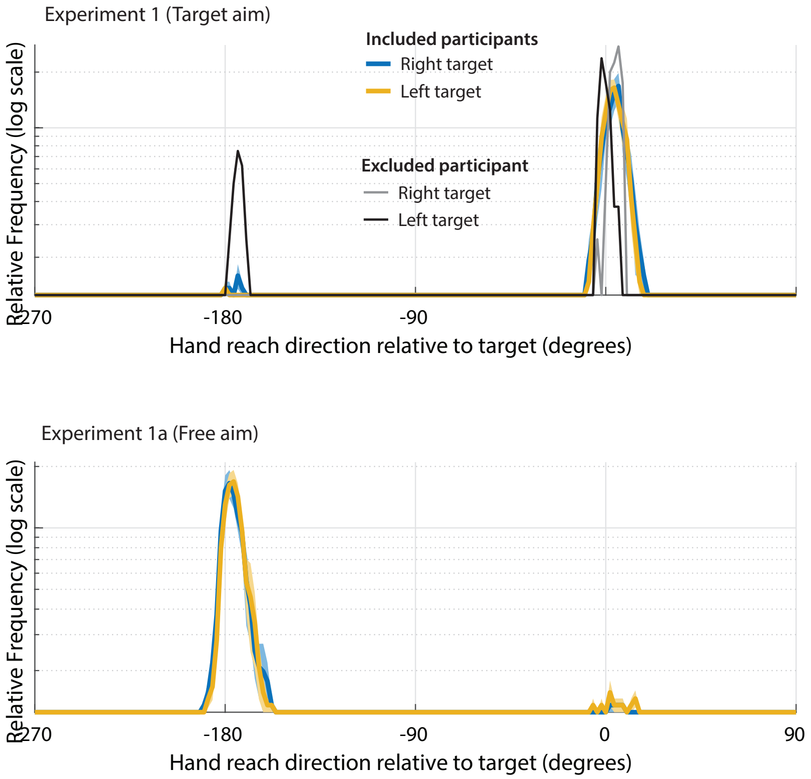


***Supplementary Figure 1: Distribution of hand reach directions for the horizontal targets in Experiments 1 and 1a.*** *In Experiment 1, reaches on the 180° direction indicate errors which suggest the aimed the cursor, rather than the hand, to the target. The (more rare) errors in Experiment 1a (where the participant reaches towards the 0deg direction) likely signify a failure to implement the strategy of aiming in the opposite direction. Note that the scale is logarithmic in order to show these relatively rare events. The thin black / gray lines show data from the Experiment 1 participant who was excluded due to a large amount of these horizontal errors (13.8% of total horizontal trials).*

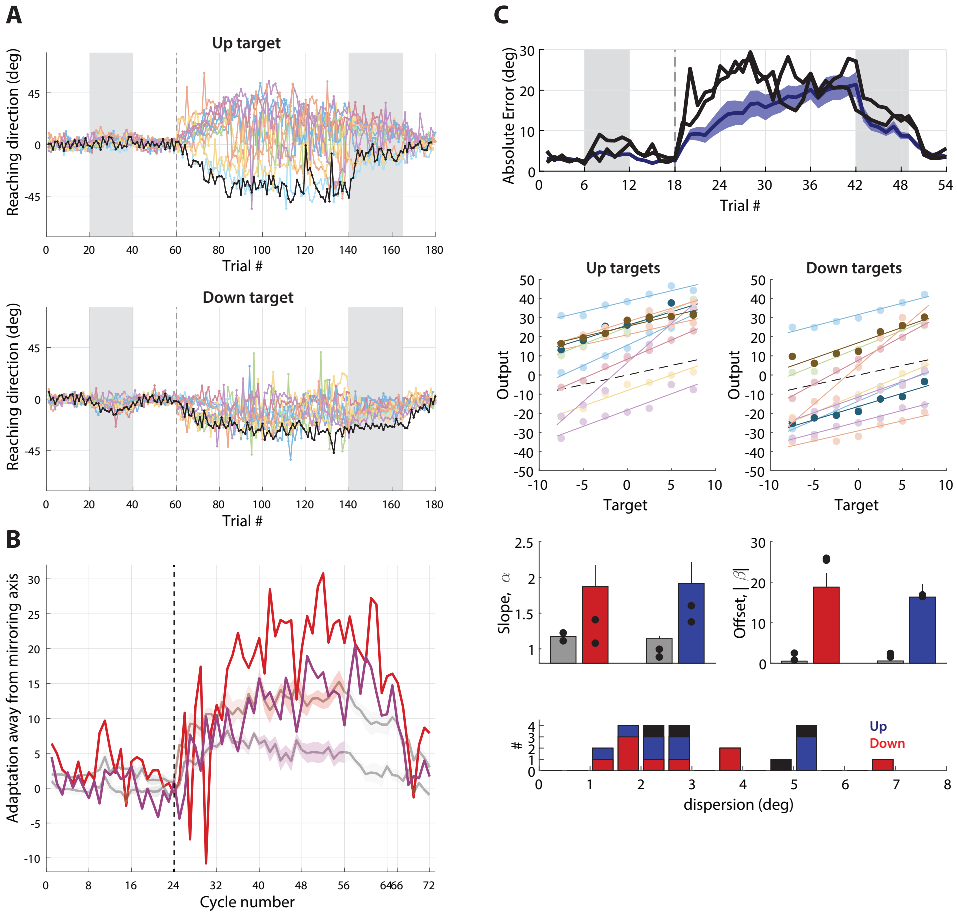


***Supplementary Figure 2: Results from participants excluded due to frequent horizontal-target errors in line with main results. A:*** *For Experiment 1, the one excluded participant (black curves) showed behavior in line with the included data (faded curves). Compare to Figure 4A. There were no participants excluded in Experiment 1a.* ***B:*** *Similarly for Experiment 2, the one excluded participant also showed adaptation away from the mirroring axis. Data overlaid against results from included participants, compare to Figure 5B.* ***C:*** *For Experiment 3, the two excluded participants, indicated by the darker curves/points, show data in line with the main results (compare to Figure 6E-H). An additional participant excluded due to erratic behavior likely due to a technical issue show in Supplementary Figure 3.*

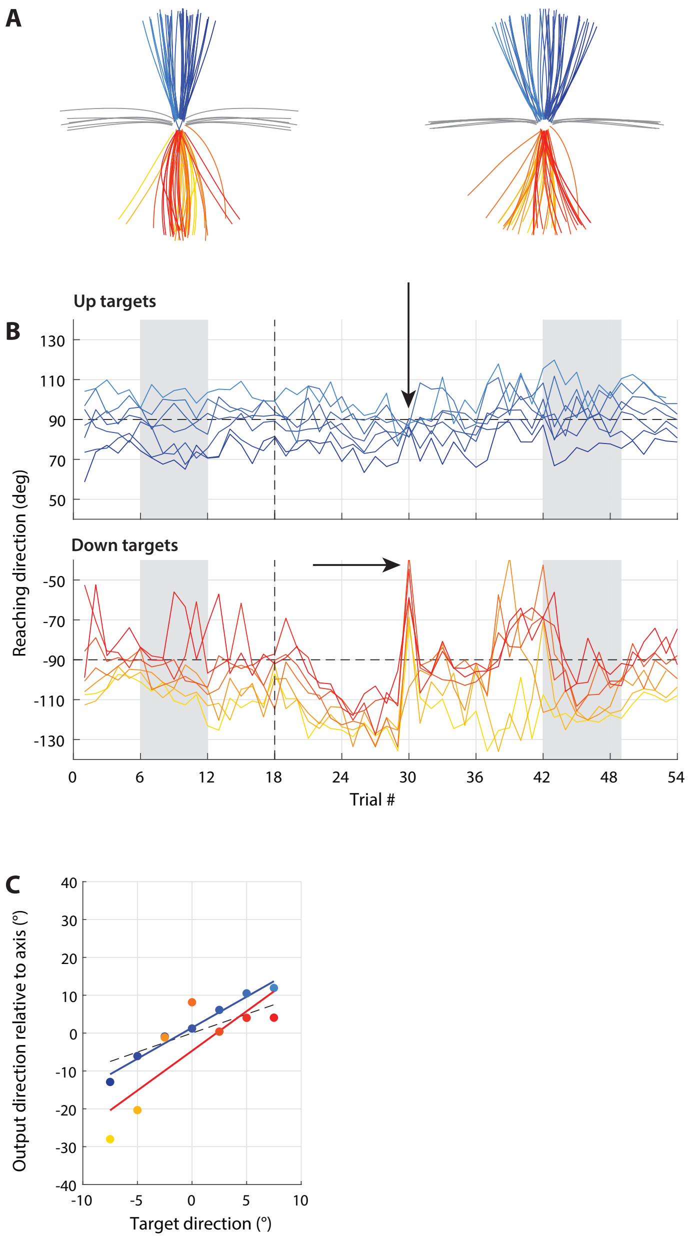


***Supplementary Figure 3:*** *Data of participant excluded due to erratic behavior in Experiment 3. Format of* ***A-C*** *as Figure 6B-D. Erratic behavior probably resulted from a large postural adjustment at the beginning of the 6*$^{th}$ *block (indicated by arrows), which would explain the abrupt changes and is in line with the participant mentioning that they had trouble reaching to the down target. Note that the result is neither in line with forward-model-based-learning (there is no reversal of the slope in panel C) nor direct-policy-update-based-learning (the target-output relationship for the down target is not strongly linear).*

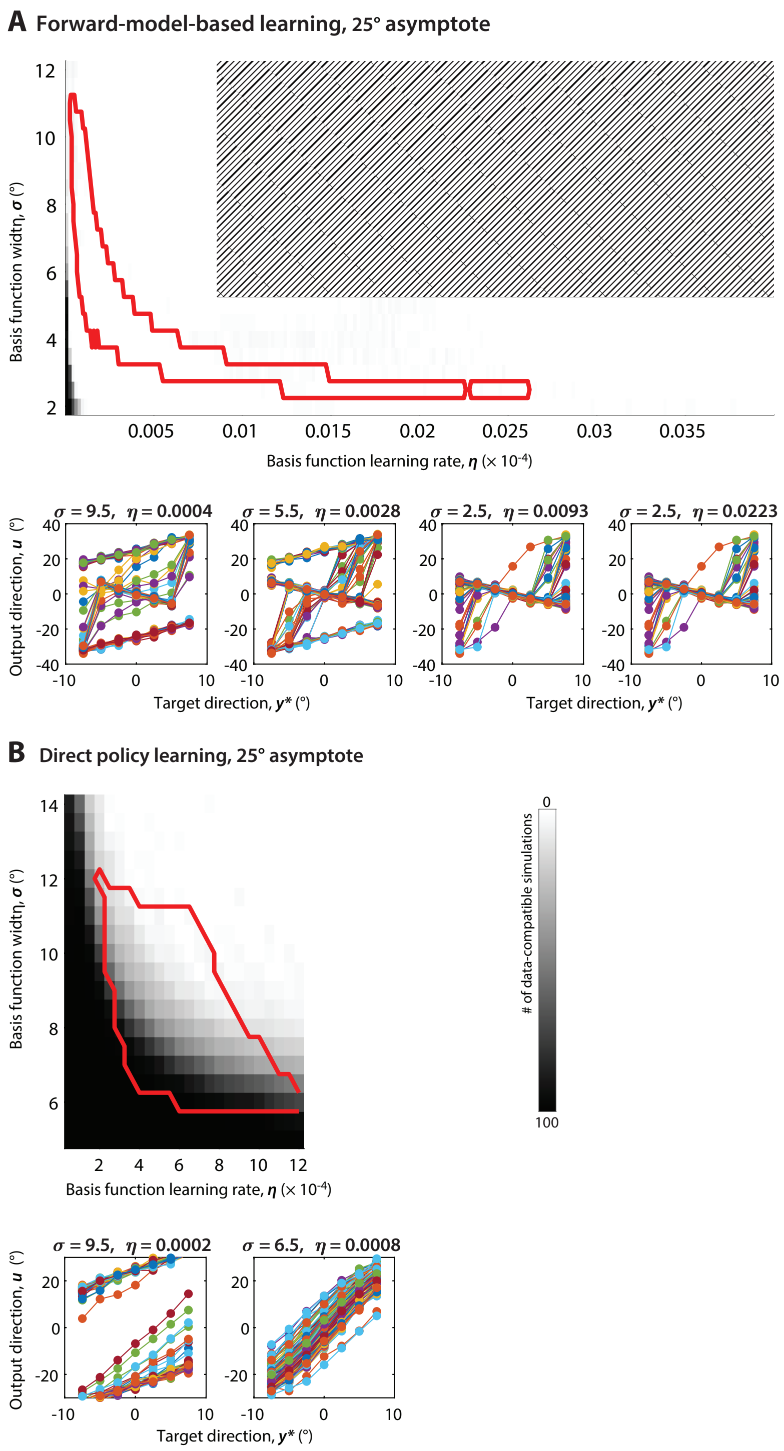


***Supplementary Figure 4:*** *Simulations of forward-model-based learning (A) and direct policy learning (B) for an asymptote of 25°. Format similar to Figure 6. As with simulations with smaller asymptotes, the data re incompatible with forward-model-based learning but compatible with direct policy learning. The additional shading in A indicates values not simulated as they were far from the range of values compatible with the learning and generalization characteristics of visuomotor adaptation in previous work (area indicated by the red line).*
